# Determining targeting specificity of nuclear-encoded organelle proteins with the self-assembling split-fluorescent protein toolkit

**DOI:** 10.1101/480962

**Authors:** Mayank Sharma, Carola Kretschmer, Christina Lampe, Johannes Stuttmann, Ralf Bernd Klösgen

**Author notes:** Mayank Sharma (Corresponding author).

## Abstract

A large number of nuclear-encoded proteins are targeted to the organelles of endosymbiotic origin, namely mitochondria and plastids. To determine the targeting specificity of these proteins, fluorescent protein tagging is a popular approach. However, ectopic expression of fluorescent protein fusions commonly results in considerable background signals and often suffers from the large size and robust folding of the reporter protein, which may perturb membrane transport. Among the alternative approaches that have been developed in recent years, the self-assembling split-fluorescent protein (*sa*split-FP) technology appears particularly promising to analyze protein targeting specificity *in vivo*. Here, we have improved this technology with respect to sensitivity and systematically evaluated its utilization to determine protein targeting to plastids and mitochondria. Furthermore, to facilitate high throughput screening of candidate proteins we have developed a *Golden Gate*-based vector toolkit, named PlaMiNGo (<u>Pla</u>stid and/or <u>Mi</u>tochondria targeted proteins <u>N</u>-terminally fused to <u>G</u>FP11 tags via *<u>Go</u>lden Gate* cloning). As a result of these improvements, dual targeting could be detected for a number of proteins, which had earlier been characterized as being targeted to a single organelle only. These results were independently confirmed with a plant phenotype complementation approach thus demonstrating the sensitivity and robustness of the *sa*split-FP-based method to analyze the targeting specificity of nuclear-encoded proteins.

**Highlight:** Several mono-specific proteins showed dual targeting to plastids and mitochondria with the self-assembling split-GFP system. A *Golden Gate*-based vector toolkit was constructed to facilitate easy cloning and subsequent determination of protein targeting specificity.

## 1. Introduction

Many nuclear-encoded proteins are targeted across membranes to reach their final destination in the cell, i.e. a sub-cellular compartment or cell organelle. To determine the specificity of such targeting, fluorescent protein tagging (FP tagging), that is, the fusion of the candidate protein with a fluorescent reporter (e.g., Green Fluorescent Protein, GFP) and subsequent *in vivo* imaging by fluorescence microscopy, is a popular approach. However, this widely utilized method carries some inherent pitfalls (reviewed by Moore and Murphy 2009). First, the large size and intrinsic folding properties of the reporter protein might perturb membrane transport of the candidate protein (Marques et al., 2004). Second, considerable background fluorescence signals due to protein overexpression and hence saturation of the organelle transport machinery is sometimes observed (Sharma et al., 2018a). And finally, proteins targeted to an organelle at low amounts are difficult to visualize using this approach due to weak fluorescence signals originating from a limited number of protein molecules.

These constraints become particularly prominent when it comes to the analysis of proteins targeted to mitochondria and/or plastids (Tanz et al., 2013). These organelle-targeted proteins usually carry an N-terminal transport signal called ‘transit peptide’ to facilitate transport of the passenger protein across the membranes via organelle specific translocation machineries. Amongst a repertoire of proteins encoded in the plant cell nucleus, more than 3000 are transported into mitochondria and/or plastids (Van Wijk and Baginsky 2011; Rao et al., 2017). While most of these proteins are targeted to only one type of organelle, a group of proteins exists with dual targeting specificity, i.e., they carry an ‘ambiguous’ transit peptide capable of translocating the passenger protein into both mitochondria and plastids (reviewed in Sharma et al., 2018b). Despite having dual targeting properties some of these proteins are still preferentially targeted to one of the two organelles. In this case, determining targeting specificity using a conventional FP-tagging approach is particularly tedious and error-prone, as highly intensive fluorescence signals coming from one organelle can mask signals of low-intensity coming from the other one (Duchêne et al., 2005, Sharma et al., 2018b).

One possibility to circumvent these problems is to detect the fluorescence signals, coming from each organelle, in separate cells. This has become technically feasible by the invention of the self-assembling green fluorescent protein (*sa*split-GFP) technology, a popular approach developed recently to determine targeting specificity of proteins *in vivo* (Cabantous et al., 2005). Spontaneous self-assembly of split-GFP relies on two highly engineered fragments derived from the ‘superfolder’ GFP variant, sfGFP. The large fragment, GFP1-10 OPT (further referred to as GFP1-10), comprises ten N-terminal antiparallel β-sheets of GFP. The smaller fragment, GFP11 M3 (further referred to as GFP11), comprises only 16 amino acids and represents the C-terminal 11^th^ β-sheet of GFP. When brought into close proximity, the two fragments can assemble spontaneously to reconstitute the functional fluorophore without the need of an additional interacting partner (Figure 1a) (Cabantous et al., 2005). For subcellular protein localization studies, GFP1-10 is fused to a transport signal of known organelle specificity and analyzed together with a chimeric protein comprising the candidate protein and GFP11. Fluorescence complementation is achieved specifically and exclusively if the transport signal of the candidate mediates transport of GFP11 into the organelle housing GFP1-10. The fluorescence signal remains limited to the compartment containing GFP1-10, irrespective of whether the GFP11-fused candidate protein is targeted to further subcellular compartments or not (Figure 1b). Thus, the *sa*split-GFP technology allows selective *in vivo* imaging of a protein of interest in a respective compartment with enhanced signal-to-noise ratio. Furthermore, the small GFP11 tag appears to have a lower propensity for interfering with membrane transport in comparison to the full length fluorescent proteins used for direct FP-tagging. Smaller size tags were previously shown to be advantageous over large FP-tags for live cell imaging (Andresen et al., 2004; Giepmans et al., 2006).

**Figure 1.**
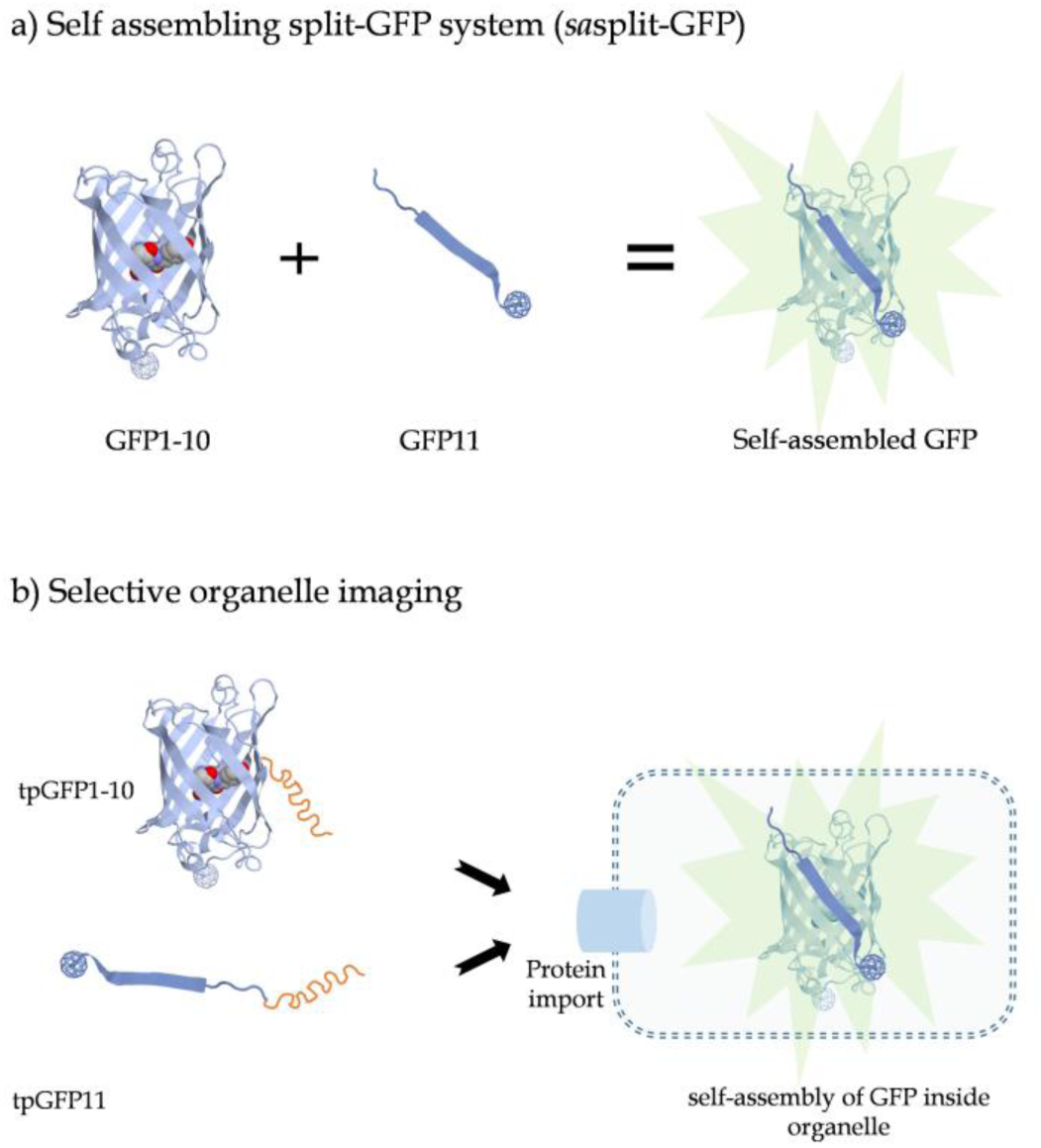
Schematic drawing illustrating the principle of the *sa*split-GFP system. **(a)** Two non-fluorescing fragments, GFP1-10 and GFP11, can self-assemble to generate a fluorescing GFP molecule. **(b)** Transport of both GFP chimeras into the same organelle is essential to achieve fluorescence signals.

The *sa*split-GFP system has already been applied to a wide range of organisms and adapted to elucidate a variety of cellular functions or processes including, for example, *in vivo* protein solubility, sub-cellular localization of pathogen effectors, endogenous protein labeling for *in vivo* imaging, protein-protein interaction studies, and membrane protein topology determination (Cabantous and Waldo 2006; Van Engelberg and Palmer, 2010; Machettira et al., 2011; Cabantous et al., 2013; Kamiyama et al., 2016; Henry et al., 2017). The principal suitability of the *sa*split-GFP system for *in vivo* imaging of protein targeting in plant cells has also been demonstrated recently (Park et al., 2017). In this case, transgenic *Arabidopsis thaliana* lines expressing the organelle-targeted GFP1-10 receptor were transiently transformed with constructs expressing a candidate protein fused to GFP11 tag. However, the requirement of transgenic ‘receptor’ plant lines and low transient transformation efficiency hampers the utilization of this approach for high throughput protein targeting studies. Along with a limited yield of transformed cells, the relatively low brightness of the *sa*split-GFP prevented the visualization of proteins targeted in low amounts into the organelles.

Therefore, we have here systematically evaluated and optimized the *sa*split-GFP system for analysis of protein targeting specificity in plant cells and assessed the effect of multimerization of the GFP11 tag on the intensity of the fluorescence signals inside mitochondria and plastids. A *Golden Gate*-based vector toolkit named PlaMiNGo (Analysis of <u>Pla</u>stid and/or <u>Mi</u>tochondrial targeted proteins <u>N</u>-terminally fused to <u>G</u>FP11 tags via *<u>Go</u>lden Gate* cloning) was developed to facilitate high-throughput analyses of candidate proteins with high transformation efficiency and enhanced signal-to-noise ratio. With this approach, dual targeting to mitochondria and plastids could be detected for several proteins that were previously characterized as being targeted to a single organelle only, which demonstrates its improved sensitivity. Importantly, plastid targeting of two of these proteins was independently confirmed using a phenotype complementation-based approach in stable transgenic *Arabidopsis* plants, which proves the *in vivo* relevance of results obtained with the PlaMiNGo system.

## 2. Results

### 2.1 The *sa*split-GFP system to determine protein targeting specificity

In order to study protein targeting into plastids, the transit peptide of a chloroplast protein, ferredoxin-NADP^+^-oxidoreductase of spinach (FNR_1-55_; Zhang et al., 2001), was fused to the GFP1-10 receptor (the large fragment of the *sa*split-GFP system) to facilitate its localization into plastids. For mitochondrial localization of GFP1-10, the N-terminal 100 amino acid residues comprising the presequence of the mitochondrial Rieske Fe/S protein of potato (mtRi_1-100_; Emmermann et al., 1994) were used. Both transport signals had previously been characterized for their targeting specificity to a single organelle with *in vivo* and *in vitro* approaches (Rödiger et al., 2011). For the initial experiments evaluating the suitability of the *sa*split-GFP system for our purposes, each transport signal was likewise combined with the GFP11_x7_ tag (small fragment of *sa*split-GFP system). A seven-fold repeat of this GFP11 tag (GFP11_x7_; separated via a 5 amino acid linker) was used, as such multiple GFP11 tags had been reported to intensify the fluorescence signals in mammalian cells (Kamiyama et al., 2016). When these constructs were co-expressed with the FNR_1-55_/GFP1-10 and mtRi_1-100_/GFP1-10 gene constructs, fluorescence signals were exclusively obtained in those instances in which the same transport signal was present in both chimeras, i.e. in plastids after co-expression of FNR_1-55_/GFP1-10 and FNR_1-55_/GFP11_x7_ and in mitochondria after co-expression of mtRi_1-100_/GFP1-10 and mtRi_1-100_/GFP11_x7_ (Figure 2). When infiltrated alone, neither of theses constructs generated any detectable fluorescence signal (Suppl. Figure 1). This demonstrates the suitability and specificity of the *sa*split-GFP system for the analysis of protein targeting.

**Figure 2.**
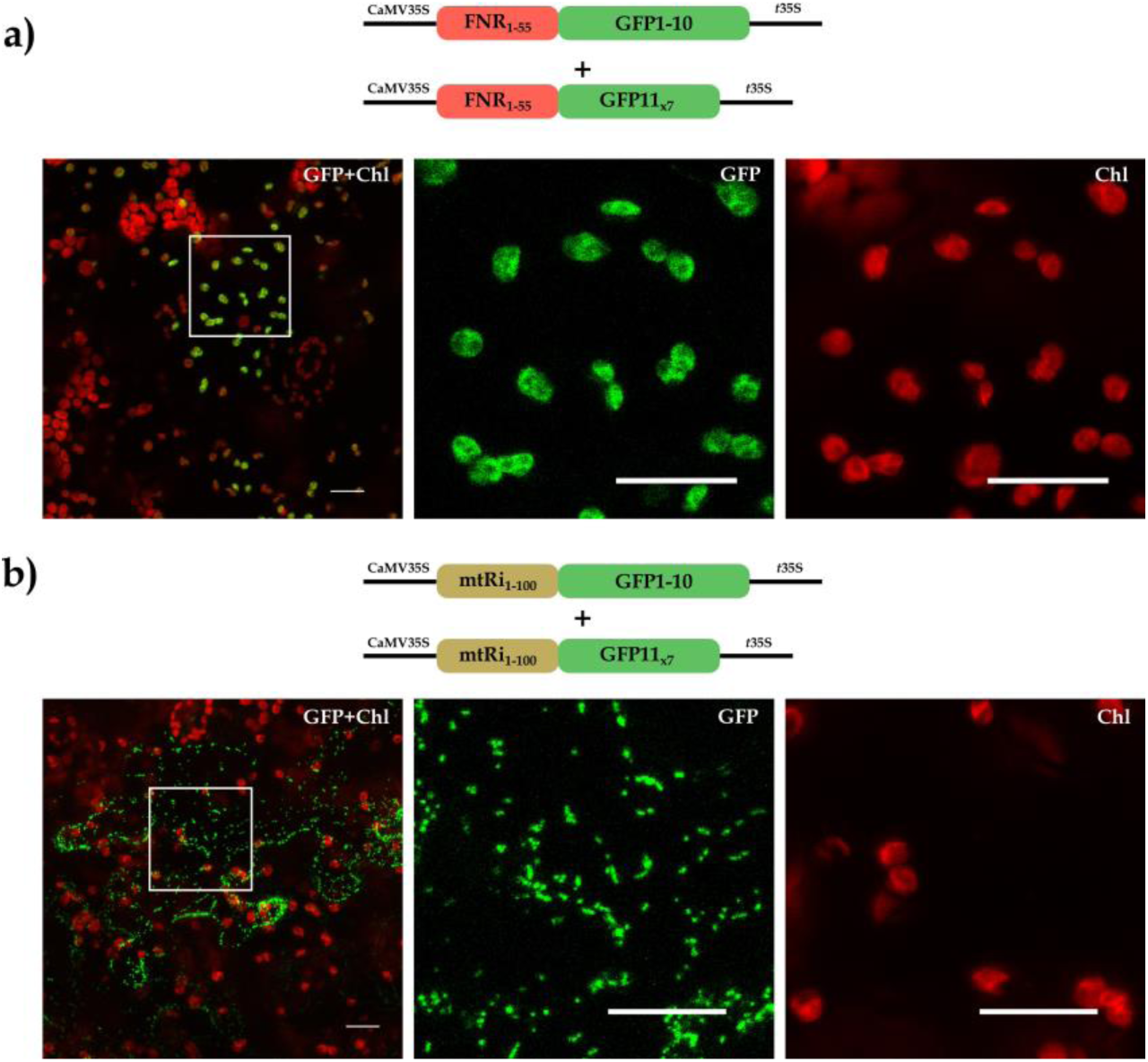
Establishment of the *sa*split-GFP system for *in vivo* organelle imaging. The coding sequences of **(a)** FNR_1-55_/GFP1-10 and FNR_1-55_/GFP11_x7_ **(b)** mtRi_1-100_/GFP1-10 and mtRi_1-100_/GFP11_x7_ were transiently co-expressed after *Agrobacterium* co-infiltration into the lower epidermis of *Nicotiana benthamiana* leaves and analyzed by confocal laser scanning microscopy (CLSM). Image acquisition of transformed cells was done with 20-x objective in several Z-stacks, which were subsequently stacked for maximum intensity projection. Representative cells (*left panels*) are presented as overlay images of the chlorophyll channel (displayed in red) and the GFP channel (displayed in green). The strong chlorophyll signals in the background are derived from the larger chloroplasts of untransformed mesophyll cells underneath the epidermal cell layers. The *squares* highlight areas of the transformed cells that are shown in higher magnification separately for the chlorophyll channel (*middle panels*) and the GFP channel (*right panel*) as indicated. Scale bars correspond to 20 μm.

Next, we wanted to evaluate the performance of this system by determining the protein targeting behavior of several dually targeted proteins. For this purpose, we have selected three previously characterized nuclear-encoded organelle proteins from *Arabidopsis thaliana* with proven dual targeting characteristics, namely TyrRS (Tyrosine-tRNA synthetase; At3g02660), GrpE (co-chaperone GrpE1; At5g55200), and PDF (peptide deformylase 1B; At5g14660) (Berglund et al., 2009; Baudisch et al., 2014). Earlier experiments employing eYFP (enhanced Yellow Fluorescent Protein) fusions had shown that all three proteins are targeted to both endosymbiotic organelles, either in comparable amounts (TyrRS_1-91_/eYFP) or preferentially to either mitochondria (GrpE_1-100_/eYFP) or chloroplasts (PDF_1-100_/eYFP) (Sharma et al., 2018a and Figure 3). The respective N-terminal amino acid sequences carrying the organelle transport signals of the candidate proteins were fused to GFP11_x7_ tags and analyzed in our system. All three candidates (TyrRS_1-91_/GFP11_x7_, GrpE_1-100_/GFP11_x7_ and PDF_1-100_/GFP11_x7_) showed targeting to both plastids and mitochondria when co-transformed with the respective organelle-targeted receptors (Figure 3). Even plastid targeting of GrpE_1-100_/GFP11_x7_ was clearly visible, which is remarkable considering the vague fluorescence signals obtained in this organelle with the ‘standard’ fluorescent reporter fusion (Figure 3b). Thus, separation of the fluorescence signals for the two organelles into different cells proved to be advantageous to determine the low plastid targeting properties of GrpE.

**Figure 3.**
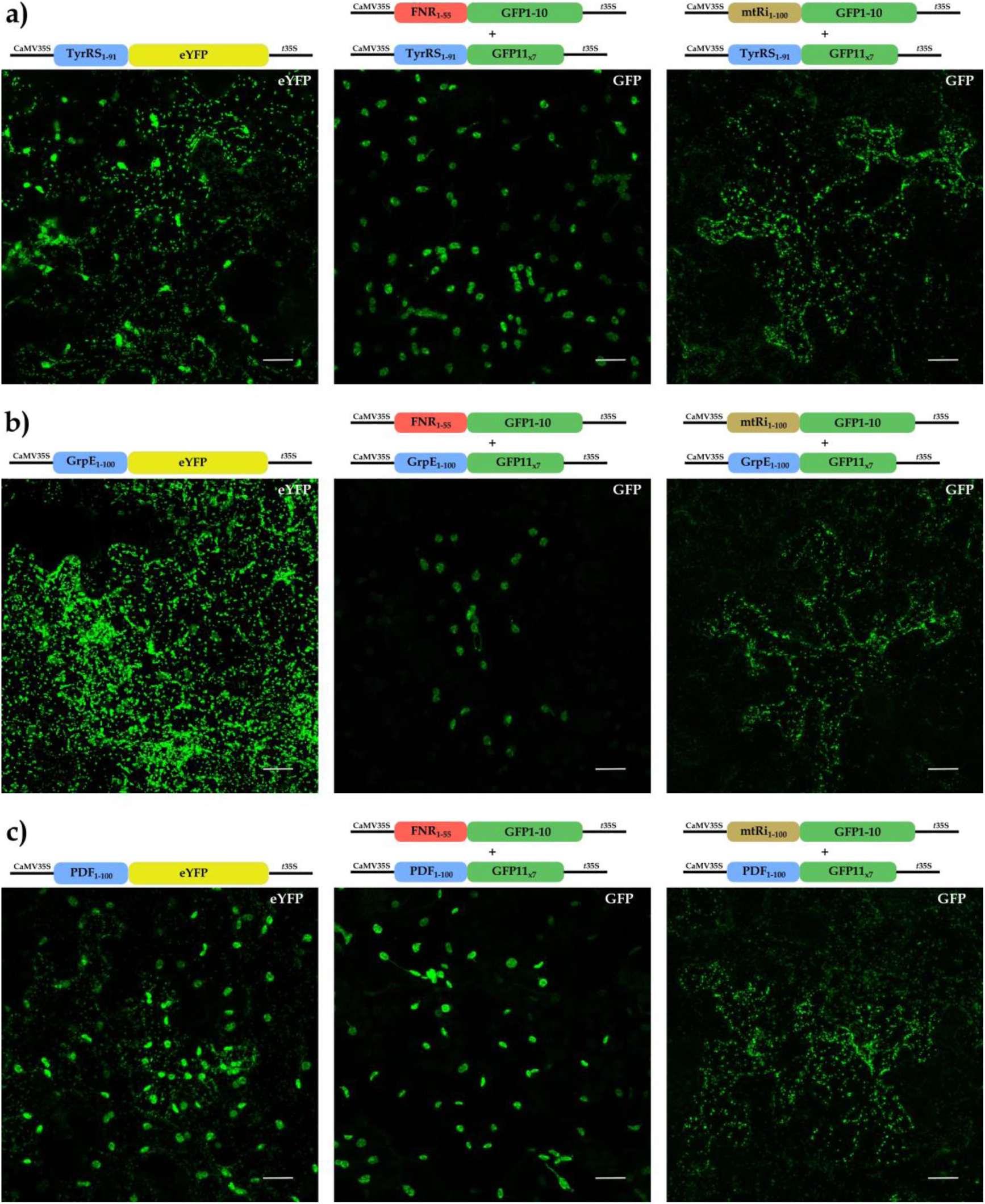
Comparison of FP-tagging and sasplit-GFP approaches to analyze the targeting specificity candidate proteins with proven dual-targeting properties. The coding sequences of **(a)** TyrRS_1-91_/GFP11_x7_, **(b)** GrpE_1-100_/GFP11_x7_ and, **(c)** PDF_1-100_/GFP11_x7_ were transiently expressed with either FNR_1-55_/GFP1-10 (*middle panels*) or mtRi_1-100_/GFP1-10 (*right panels*) via *Agrobacterium* co-infiltration into the lower epidermis of *Nicotiana benthamiana* leaves and analyzed by CLSM. For comparison the respective eYFP fusions namely **(a)** TyrRS_1-91_/eYFP, **(b)** GrpE_1-100_/eYFP, and **(c)** PDF_1-100_/eYFP were transiently expressed as well and analyzed by CLSM (*left panels*). For further details see the legend of Figure 2. Scale bars correspond to 20 μm.

### 2.2. Multimerization of the GFP11 tag leads to fluorescence signal enhancement in plastids but not in mitochondria

In the initial experiments described above, we have used GFP11_x7_, the seven-fold repeat of the GFP11 tag. However, the requirement or benefits of such multiple GFP11 tags to enhance fluorescence signals in mitochondria and plastids of plant cells were not systematically assessed. Hence, we have compared the fluorescence signal intensity obtained in the two organelles with either seven (GFP11_x7_), three (GFP11_x3_) or a single repeat (GFP11_x1_) of the GFP11 tag when fused to the dually targeted TyrRS_1-91_ peptide as protein transport signal. It turned out that the use of a single GFP11 tag yields only very faint signals in plastids while the GFP11_x3_ and GFP11_x7_ tags significantly enhance the fluorescence signals (Figure 4a) supporting the assumption that multiple GFP11 repeats can boost fluorescence signal intensity in this organelle. In contrast, for mitochondria no such correlation of number of GFP11 tag repeats and fluorescence signal intensity was found. Instead, the signal intensities were largely similar for all three constructs (Figure 4b). However, since the fluorescence signals obtained with the single GFP11 tag in mitochondria were brighter than in plastids, they are usually sufficient for proper visualization of organelle.

**Figure 4.**
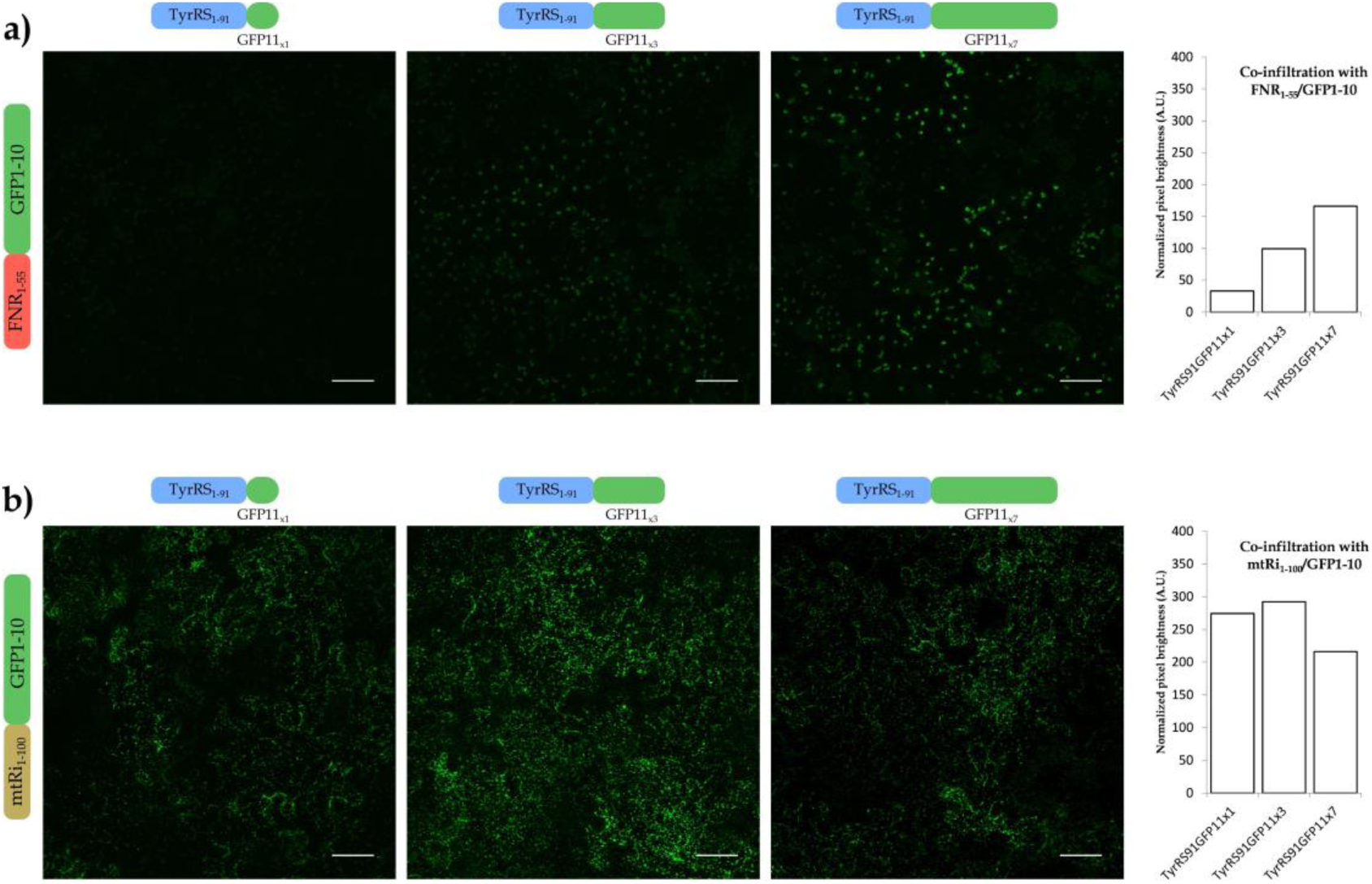
Effect of multiple GFP11 tags on fluorescence signal intensity. The gene coding sequence of the dual targeting transit peptide TyrRS_1-91_ was fused with different GFP11 variants and co-expressed with either **(a)** FNR_1-55_/GFP1-10 or **(b)** mtRi_1-100_/GFP1-10 in *Nicotiana benthamiana* lower epidermal cells. The graph represents normalized average pixel brightness obtained after subtraction of background fluorescence signals in arbitrary units (A.U.) from 9 maximum intensity projected images for each experiment, obtained from two plant replicates (see Suppl. Figure 2 for absolute quantification). Scale bars correspond to 50 μm.

### 2.3 Construction of the PlaMiNGo toolkit

To facilitate easy combination of the *sa*split-GFP technology and fluorescence signal enhancement for high-throughput screening of protein targeting specificity, we have developed a set of *Golden Gate*-based vectors. Destined to analyze the targeting specificity of candidate proteins to plastids and mitochondria, these vectors facilitate easy cloning of the candidate proteins upstream of single or multiple GFP11 tags. In this PlaMiNGo toolkit, we have utilized highly efficient gene regulatory elements, namely a ‘long’ 35S promoter (Engler et al., 2014) and *ocs* or *rbcs E9* transcriptional terminators (De Greve et al., 1982; Coruzzi et al., 1984), to control the expression of the chimeric genes, (Figure 5a and Suppl. Figure 3). Moreover, to avoid the requirement for performing co-transformation using two plasmids, we have utilized a single T-DNA expression system (Grefen and Blatt, 2012; Hecker et al., 2015) comprising two gene expression cassettes. This has the advantage that each transformed cell expresses both chimeras simultaneously. Consequently, the final vectors contain expression cassettes, for GFP1-10 gene chimeras to be targeted to either plastid or mitochondria by fusion with mtRi_1-100_ or FNR_1-55_ and another for one, three or seven times repeat of GFP11 tags. The latter expression cassette furthermore carries a *ccd*B cassette upstream of the GFP11 tag, which can be replaced by the candidate gene in a single *Golden Gate* reaction step allowing for background-free selection of positive clones. As a result, six vectors, three destined to analyze potential plastid targeting and three for mitochondria targeting analysis, were generated (Figure 5a and Suppl. Figure 3).

**Figure 5.**
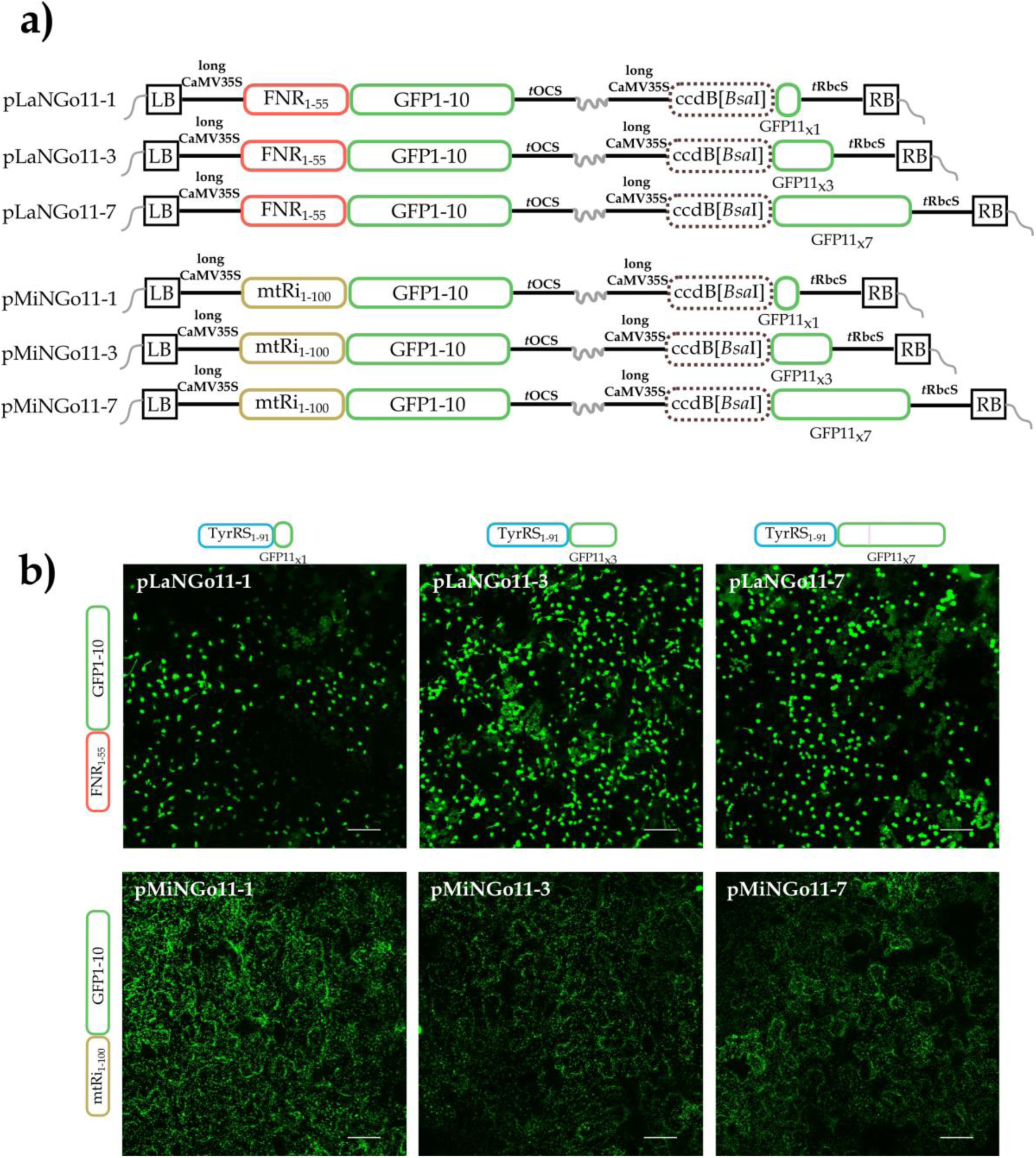
PlaMiNGo toolkit **(a)** Schematic representation of the vectors generated (see Supplementary Figure 3 for details) **(b)** PlaMiNGo vectors carrying the dually targeted TyrRS_1-19_ transit peptide at the N-terminus of GFP11 tags were used to transform *Nicotiana benthamiana* lower epidermis cells via *Agrobacterium* infiltration. The quantification of signals is presented in supplementary Figure 2. Scale bars correspond to 50 μm.

To evaluate the functionality of these vectors, the dual targeting transport signal of TyrRS (N terminal 1-91 amino acids) was cloned upstream of the GFP11 tags of all six vectors and analyzed via *Agrobacterium*-mediated transient transformation of *Nicotiana benthamiana* leaves. In all instances, tremendously improved signal intensities were observed for both organelles (Figure 5b and Suppl. Figure 2). Now, even in plastids a single copy of the GFP11 tag was sufficient for reliable detection of the reconstituted GFP fluorescence, although considerable signal enhancement could still be observed with GFP11_x3_ and GFP11_x7_ tags. This suggests that the GFP11_x7_ tag, when co-expressed with the plastid targeted GFP1-10 receptor, should allow for detection even of minute amounts of plastid-localized proteins. Likewise, upon imaging of mitochondria the fluorescence signals obtained with the single GFP11 tag were more than 3-fold stronger than the signals obtained in the previous experiments using separate vectors. Still, as observed earlier, no further improvement of signal intensities by multimerization of the GFP11 tag could be obtained in mitochondria (Suppl. Figure 2b). Instead, artificial protein aggregates were observed in some cells expressing constructs with multiple GFP11 tags (GFP11_x3_ and GFP11_x7_) (Suppl. Video 1).

### 2.4 Multicolor imaging of dual protein targeting to two organelles

A further objective of the study was to establish simultaneous multicolor imaging with the *sa*split-FP system. For this purpose, we have modified the GFP1-10 receptor to generate a yellow shifted variant (YFP1-10) using a single amino acid substitution (T203Y) as reported earlier (Kamiyama et al., 2016). To test if this variant can assemble with GFP11 to generate a functional fluorophore inside the organelles, the plastid (FNR_1-55_) or mitochondria (mtRi_1-100_) transport signals were separately fused to the N-terminus of YFP1-10. These fusions were co-infiltrated with a gene construct encoding the dual targeting transport signal TyrRS_1-91_ fused to either GFP11_x1_ or GFP11_x7_. When imaged with an YFP-specific filter-set, fluorescence signals were solely obtained in mitochondria (with mtRi_1-100_/YFP1-10 and TyrRS_1-91_/GFP11_x1_) or in plastids (with FNR_1-55_/YFP1-10 and TyrRS_1-91_/GFP11_x7_), demonstrating that the YFP1-10 fragment can indeed assemble with GFP11 in both organelles (Figure 6a, b).

**Figure 6.**
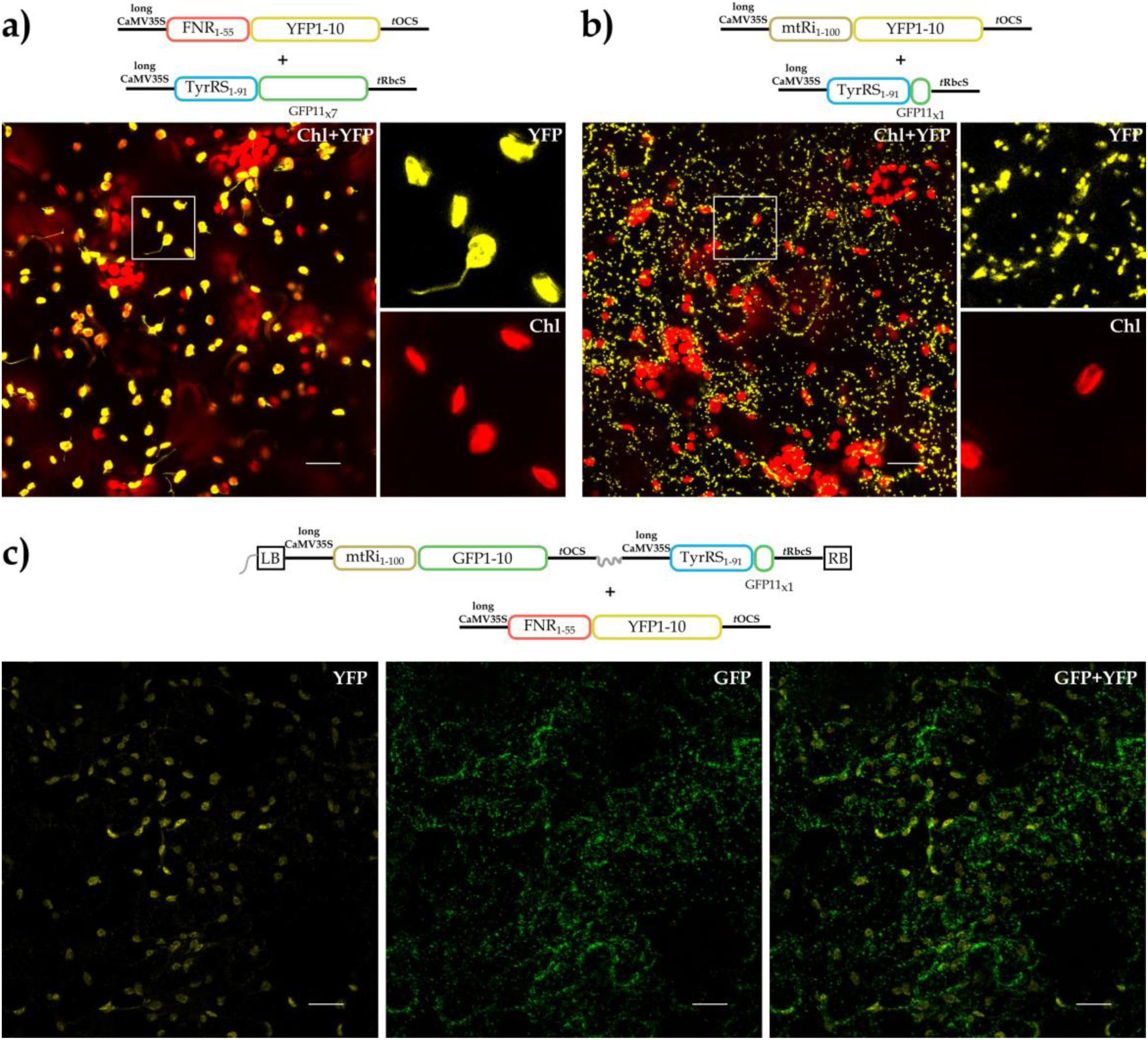
Multicolor imaging with *sa*split-YFP. The gene coding sequences of **(a)** FNR_1-55_/YFP1-10 and TyrRS_1-91_/GFP11_x7_, or **(b)** mtRi_1-100_/YFP1-10 and TyrRS_1-91_/GFP11_x1_ were transiently co-expressed in *Nicotiana benthamiana* leaf epidermis and analyzed by CLSM. **(c)** For the purpose of simultaneous multicolor imaging within the same cell, the PlaMiNGO vector comprising mtRi_1-100_/GFP1-10 and TyrRS_1-91_/GFP11_x1_ was co-infiltrated with FNR_1-55_/YFP1-10 resulting in self-assembly of GFP in mitochondria and YFP in plastids. Scale bars correspond to 20 μm.

Next, we have tested if multicolor imaging, i.e. the simultaneous labeling of plastids and mitochondria with YFP1-10 and GFP1-10, respectively, within the same cell is possible. For this purpose, the FNR_1-55_/YFP1-10 fusion was co-infiltrated with a vector comprising mtRi_1-100_/GFP1-10 and TyrRS_1-91_/GFP11_x1_. This should result in transformed cells co-expressing the dually targeted GFP11 (via fusion with TyrRS_1-91_) and two different receptors targeted to two different organelles, namely YFP1-10 to plastids (FNR_1-55_/YFP1-10) and GFP1-10 to mitochondria (mtRi_1-100_/GFP1-10). Indeed, the resulting transformed cells emitted fluorescence signals of different spectra in the two organelles due to reassembly of the GFP11 tag with both, the GFP1-10 and YFP1-10 receptors within mitochondria and plastids, respectively (Figure 6c). However, in all transformed cells a certain degree of “bleed through” of signals, i.e., the appearance of YFP fluorescence signals in GFP channel and *vice versa,* could be detected. The adjustment of the filter-sets to avoid such “bleed through” inevitably led to significant reduction of the fluorescence signal intensity. In summary, the YFP1-10 derivative of *sa*split-GFP system is not yet perfect for multicolor imaging but represents a promising basis for development of such tools.

### 2.5 *Analysis* of protein targeting specificity with PlaMiNGo

Finally, we have examined the suitability of the PlaMiNGo toolkit using eight candidates with presumed targeting specificity either to plastids, mitochondria, or to both organelles (Table 1). Three of these proteins, namely Gtred (Monothiol glutaredoxin-S15), GCS (Glycine cleavage system H protein 1) and GAPDH (Glyceraldehyde-3-phosphate dehydrogenase B) had earlier been reported to be dually targeted (Baudisch et al., 2014). Two other candidates, namely FNR and RbcS (small subunit of Rubisco from pea), are well-characterized plastid proteins (Highfield and Ellis 1978; Zhang et al., 2001), while the residual three candidates, namely mtRi, ATPS (ATP synthase subunit beta-3) and CoxIV (Cytochrome *c* oxidase subunit IV) of yeast, are known for their mitochondrial targeting specificity (Maarse et al., 1984; Emmermann et al., 1994; Baudisch et al., 2014). The protein fragments comprising the transport signals of these proteins were cloned as fusions with GFP11 tags into the PlaMiNGo vectors, additionally comprising the organelle targeted receptor fusions FNR_1-55_/YFP1-10 or mtRi_1-100_/YFP1-10.

**Table 1.** Sub-cellular localization of candidate proteins as determined with different approaches.

| Candidate | *sasplit*-GFP | *in vitro* import<sup>¶</sup> | FP Localization | Gene Accession |
| --- | --- | --- | --- | --- |
| FNR | Chloro | Chloro<sup>2</sup> | Chloro<sup>2</sup> | M86349.1 |
| mtRi | Mito | Mito<sup>2</sup> | Mito<sup>2</sup> | X79332.1 |
| GCS | Mito and Chloro | Mito and Chloro<sup>2</sup> | Mito and Chloro<sup>2</sup> | At2g35370 |
| GAPDH | Mito and Chloro | Mito and Chloro<sup>2</sup> | Chloro<sup>2</sup> | At1g42970 |
| Gtred | Mito and Chloro | Mito and Chloro<sup>2</sup> | Mito and Chloro<sup>2</sup>, Chloro<sup>4</sup>, Mito<sup>5</sup> | At3g15660 |
| CoxIV | Mito and Chloro | NA | Mito<sup>6</sup> | SGD:S000003155 |
| ATPS | Mito and Chloro | Mito<sup>2</sup> | Mito<sup>2</sup> | At5g08680 |
| SSU | Mito and Chloro | Mito and Chloro<sup>1</sup> | Chloro<sup>3</sup> | XM_016585367 |

<sup>¶</sup>Import studies were performed with the authentic precursor proteins; NA- not available ; <sup>1</sup>Rudhe et al., 2002; <sup>2</sup>Baudisch et al., 2014; <sup>3</sup>Nelson et al., 2007; <sup>4</sup>Cheng, 2008; <sup>5</sup>Moseler et al., 2015; <sup>6</sup>Köhler et al., 2003; *Mito*- mitochondria; *Chloro*- Chloroplasts or plastids.

Five of the eight candidates showed in our assay system the same targeting behavior as reported in the literature (Table 1): mtRi and FNR showed exclusive transport into either mitochondria or plastids, respectively (Figure 7a-b) (Rodiger et al., 2011), and the dual targeting candidates GCS, Gtret and GAPDH showed transport into both organelles (Baudisch et al., 2014) (Figure 7c-e). It should be noted though that in the case of GAPDH mitochondrial targeting was rather weak in our assays and could be observed only in few transformed cells. However, it was rather unexpected that the remaining three monospecific candidates, namely ATPS_1-100_, RbcS_1-79_ and CoxIV_1-29_, showed dual targeting in our experiments (Figure 8). In the literature, mitochondrial targeting of RbcS has already been described once (Rudhe et al., 2002) but these results were solely based on *in vitro* assays. In addition, dual targeting of the yeast mitochondria presequence, CoxIV, could be assumed considering high degree of freedom of non-plant mitochondria transport signals (Staiger et al., 2009). However, this dual targeting was entirely unexpected for the plant mitochondrial protein, ATPS_1-100_, which never showed any targeting to plastids when analyzed with *in vivo* fluorescent protein tagging and *in vitro* protein transport experiments (Baudisch et al., 2014 and Suppl. Figure 4).

**Figure 7.**
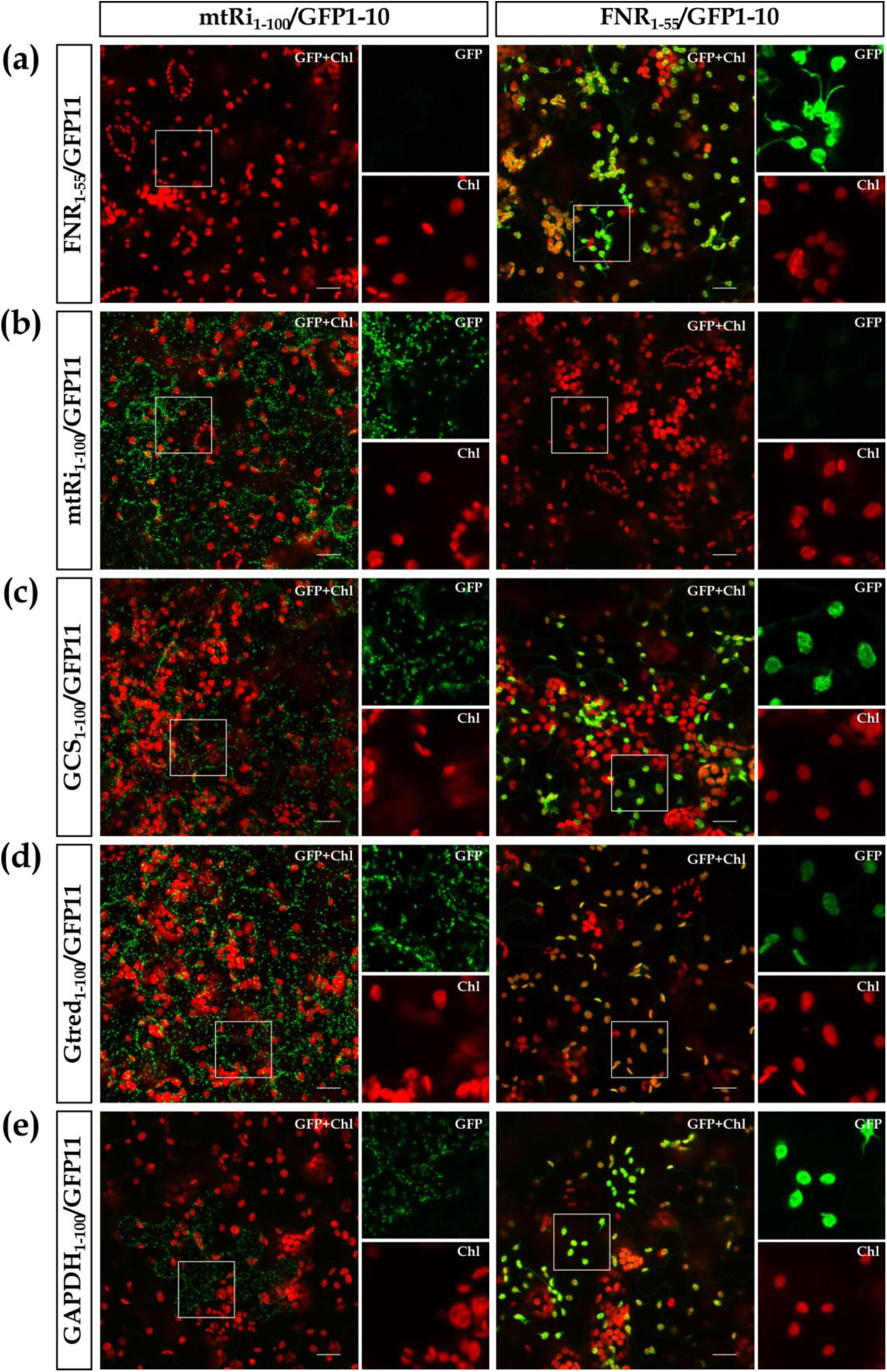
Analysis of candidate proteins with the PLaMiNGo toolkit. Gene fragments encoding the transport signals of either **(a)** FNR, **(b)** mtRi, **(c)** GCS, **(d)** Gtred or **(e)** GAPDH, were cloned upstream of GFP11_x1_ and GFP11_x7_ tags in the two PlaMiNGo vectors comprising either mtRi_1-100_/GFP1-10 (*left panels*) or FNR_1-55_/GFP1-10 (*right panels*) respectively. The resulting constructs were used to transform the lower epidermis of *Nicotiana benthamiana* leaves via *Agrobacterium* infiltration. For further details see the legend of Figure 2. Scale bars correspond to 20 μm.

**Figure 8.**
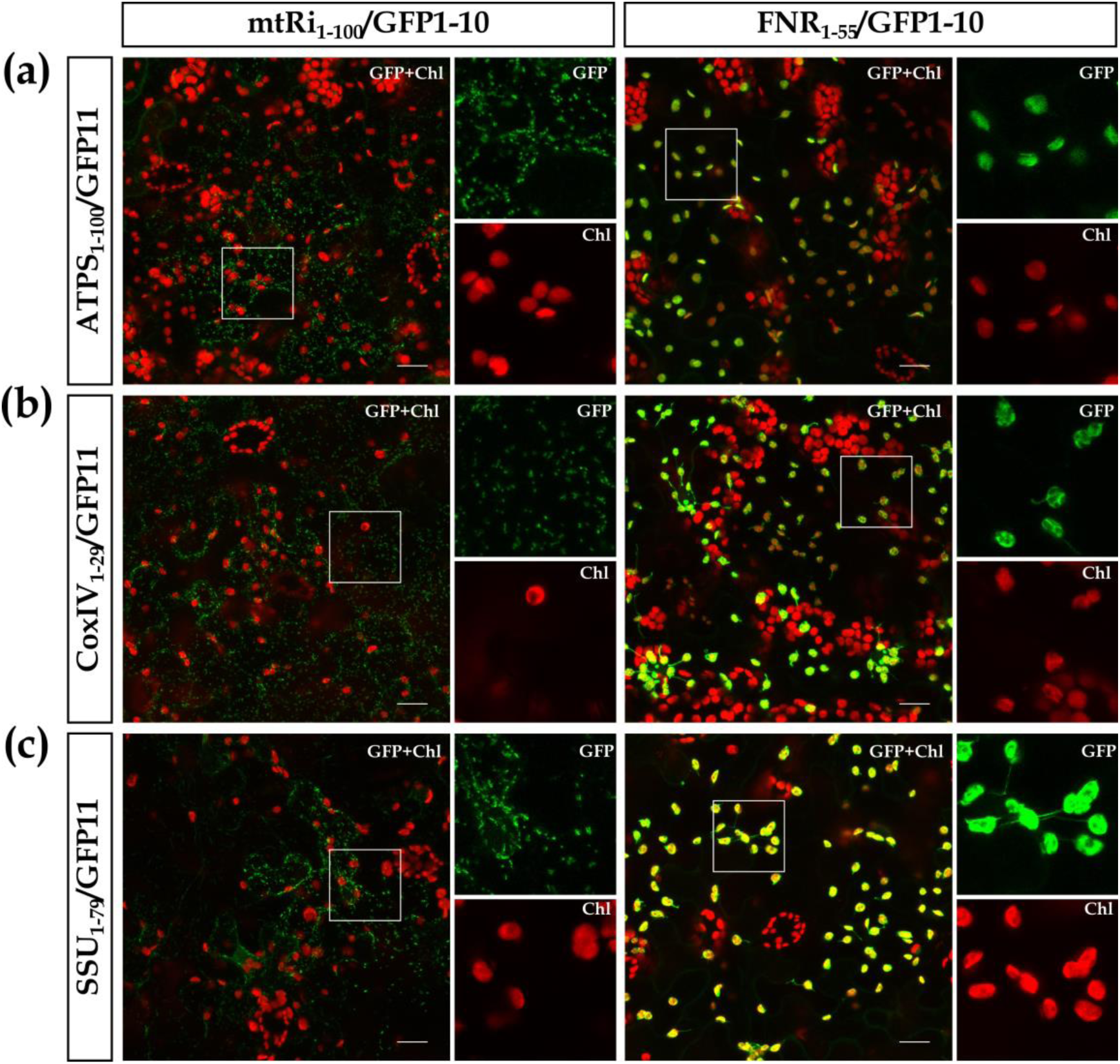
Dual localization of presumed monospecific candidate proteins. Gene fragments encoding the transport signals of either **(a)** ATPS, **(b)** CoxIV or **(c)** SSU, were cloned upstream of the GFP11_x7_ tag in the two PlaMiNGo vectors comprising either mtRi_1-100_/GFP1-10 (*left panels*) or FNR_1-55_/GFP1-10 (*right panels*). The resulting constructs were used to transform the lower epidermis of *Nicotiana benthamiana* leaves via *Agrobacterium* infiltration. For further details, see the legend of Figure 2. Scale bars correspond to 20 μm.

### 2.6 Phenotype complementation confirms the plastid targeting properties of ATPS

These unexpected results demanded for independent confirmation. Thus, to re-evaluate the plastid targeting properties of the transport signal of mitochondrial ATPS we have applied a phenotype complementation approach. In this approach, we made use of the *immutans* mutant of *Arabidopsis thaliana* that shows a white-green sectored (variegated) leaf phenotype and stunted growth when grown under daylight conditions. The mutant phenotype is due to the absence of a functional nuclear-encoded plastid protein, namely plastid terminal oxidase (PTOX; Carol et al., 1999; Aluru et al., 2001). Complementation of the variegated phenotype of *immutans* requires a functional PTOX protein in plastids (Fu et al., 2005 and Figure 9c). To adapt PTOX for our analysis, we have replaced the authentic transit peptide of PTOX with the N-terminal 100 AA residues of two candidate proteins, namely ATPS and GCS (a validated dually targeted protein) generating ATPS_1-100_/mPTOX and GCS_1-100_/mPTOX, respectively. Two further constructions encoding either the authentic PTOX precursor or the transit peptide-free mature PTOX protein (mPTOX) were generated for comparison. These gene chimeras were expressed under the control of the CaMV35S promoter in the *immutans* mutant plants. As expected, both the authentic PTOX precursor and GCS_1-100_/mPTOX were able to complement the variegated phenotype, as expected. In contrast, the expression of mPTOX cannot complement the mutant phenotype (Figure 9). These results confirm that targeting of PTOX into plastids is essential for complementation of the *immutans* variegated phenotype. Remarkably, also the expression of ATPS_1-100_/mPTOX resulted in mutant phenotype complementation (Figure 9f). However, such complementation was observed in only four from six transgenic lines analyzed suggesting that plastid targeting of ATPS_1-100_/mPTOX is, in principle, possible but apparently less efficient than with typical plastid targeted transit peptides. These results clearly underline the basic plastid targeting properties of mitochondrial protein ATPS and thus re-confirm the results obtained with the *sa*split-GFP technology established here.

**Figure 9.**
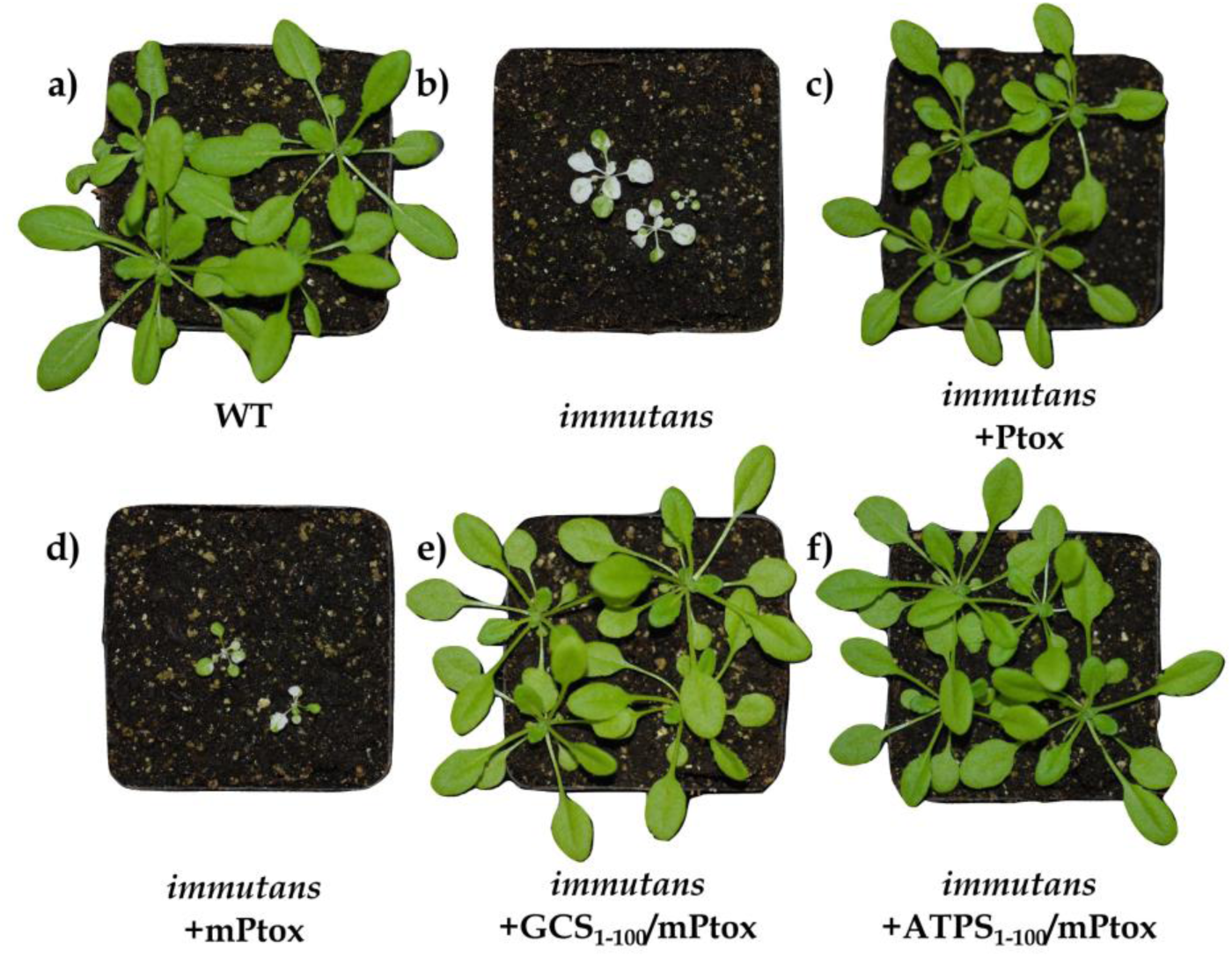
Complementation analysis of the *immutans* mutant of *Arabidopsis thaliana*. Phenotype of **(a)** wild type Col-0 and **(b)** *immutans* mutant plants. Gene constructions encoding either **(c)** the precursor PTOX (Ptox), **(d)** the mature PTOX (mPTOX) or PTOX fused to the transit peptides of the candidate proteins **(e)** GCS or **(f)** ATPS, were used to transform *immutans* mutant plants for generation of complementation lines. At least three independently transgenic lines were examined for variegated phenotype complementation of the variegated *immunats* phenotype. The figure shows representative 5 weeks old T2 generation plants.

## 3. Discussion

The goal of this study was to assess the suitability of the signal enhanced *sa*split-GFP system to determine the targeting specificity of nuclear-encoded organelle proteins and to develop tools for rapid cloning and subsequent analysis of their targeting behavior. The characterization of the targeting specificity of organelle proteins is crucial (i) to elucidate the functional properties of the organelle transport machineries, (ii) to study evolutionary aspects of protein targeting, and (iii) for biotechnological applications employing organelles. The application of the *sa*split-GFP system, as demonstrated in this study, provides a novel toolbox to quickly determine the targeting properties of candidate proteins with high sensitivity.

### 3.1 Selective imaging and fluorescence signal enhancement with *sa*split-GFP technology

#### 3.1.1 Selective Imaging

The selective imaging of organelles is one of the major advantages of the *sa*split-GFP system in comparison to ‘standard’ fluorescent protein tagging approaches. The requirement for the presence of the non-fluorescing GFP1-10 receptor in a specific subcellular location is the key for selective imaging (Kaddoum et al., 2010). In this study, two transport signals, namely FNR_1-55_ and mtRi_1-100_, were selected for localization of the receptor specifically within two sub-cellular locations, the plastid stroma and the mitochondrial matrix, respectively. As a result, fluorescence signals will appear only if the GFP11 tagged protein is completely imported into the same sub-cellular location and not if a protein is merely binding to the organelle surface. This was otherwise difficult to distinguish with FP-based approaches, specifically for mitochondria due to their small size.

#### 3.1.2 Fluorescence signal enhancement

The self-assembling split-GFP molecules have been reported to produce fluorescent signals of lower intensity than ‘standard’ fluorescent proteins (Kökker et al., 2018). This problem can be circumvented with the use of multiple GFP11 tags. However, the two organelles respond differently to this modification. While signal enhancement with multiple GFP11 tags works well in plastids, fluorescence signal enhancement could not be observed in mitochondria (Figure 4). One possible reason might be the size difference between these organelles. Plastids are much larger in size and thus the proteins in the plastid stroma are present in lower concentration. Consequently, the chances for self-assembly of *sa*split-GFP fragments in this organelle are lower than in mitochondria and thus fluorescence signal enhancement could be observed by increasing number of GFP11 tags in plastids. Furthermore, differences in the physicochemical properties of the two organelles, e.g. pH, might likewise contribute to this phenomenon.

The use of more efficient gene regulatory elements, i.e. promoter and terminator, in the PlaMiNGo toolkit also led to significant enhancement in fluorescence signal intensity. However, in combination with multiple GFP11 tags such increased gene expression can lead to the formation of aggregations in transformed cells, particularly if these tags are combined with mitochondria targeting transport signals. The intrinsic property of the *sa*split-GFP fragments to form dimers and aggregates (Cabantous et al., 2005) and comparatively less efficient unfoldase activity of the mitochondrial protein translocation machinery (Agarraberes and Dice 2001) could be one of the possible reasons for this phenomenon. Since, the protein unfolding prior to translocation is apparently more efficient in plastids, the multiple GFP11 tags can efficiently be imported into this organelle. On the other hand, high expression of plastid targeted GFP1-10 alone could result in appearance of faint fluorescence signals, probably due to formation of dimers. In most instances, these faint signals are clearly distinguishable from ‘actual’ fluorescence signals obtained via self-assembly of split-GFP but still it should considered as an experimental control to avoid the misinterpretation of results

### 3.2 High sensitivity of *sa*split-fluorescence protein system

Fluorescent signal enhancement in combination with selective imaging makes the *sa*split-GFP system highly sensitive with respect to targeting specificity determination. In consequence, dual targeting of several proteins was newly detected with the PlaMiNGo toolkit developed here, which had previously been missed due to inherent limitations of ‘standard’ fluorescent protein tagging approaches. For example, GAPDH shows dual targeting with the *sa*split-GFP system, in line with the results of *in vitro* import experiments (Baudisch et al., 2014). In contrast, with ‘classical’ *in vivo* approaches using FP-tagging, GAPDH appeared to be solely transported into plastids (Baudisch et al., 2014). Similarly, as shown here, the transit peptide of RbcS is able to translocate the GFP11 tag into mitochondria, but this property remained undetected with the FP-tagging approach. Remarkably, such mitochondria targeting properties of the RbcS transit peptide were also found in a recent study employing sulfadiazine-resistant plants (Tabatabaei et al., 2018). In contrast, dual targeting of the transport signal of yeast CoxIV had never been reported earlier. Dual targeting of yeast mitochondrial transport signal might be a consequence of the fact that yeast does not contain plastids and thus the transport signal of yeast mitochondria had not ‘learned’ to distinguish between the two endosymbiotic organelles (Staiger et al., 2009). The plastid targeting of a yeast mitochondria transport signal has also been reported earlier (Huang et al., 1990). However, such dual targeting was most unexpected for the transport signal of ATPS because neither *in vitro* nor *in vivo* approaches gave any hint of the plastid targeting properties of this transport signal (Baudisch et al., 2014). Even transgenic plants expressing ATPS_1-100_/eYFP did not show any plastid localization (Suppl. Fig. 4). The fact, that it was clearly detectable here and that this result could be independently confirmed by a phenotype complementation approach using *immutans* mutants (Figure 8a, 9f), underlines the sensitivity of the *sa*split-GFP technology.

### 3.3 Modularity of PlaMiNGO toolkit

The vector toolkit is based on the principle of modular cloning (Weber et al., 2011). Hence, the components of the PlaMiNGo toolkit can easily be rearranged for *in vivo* imaging of proteins targeted to various other sub-cellular compartments. The vectors constructed in module 1 (Suppl. Figure 3) are binary vectors and can be utilized for plant cell transformation via *Agrobacterium* or via several other methods, e.g. protoplast transformation or particle bombardment. The gene of interest can be cloned upstream to one of the GFP11 tags with a single *Golden Gate* cloning reaction. Similarly, GFP1-10 can be targeted to the different sub-cellular or even sub-organellar compartments via cloning of the specific transport signal N-terminally in a *Golden Gate ‘*ready’ vector (pTEI176 or pTEI177). These vectors carry a *ccd*B negative selection cassette upstream of GFP1-10 with *Bsa*I restriction sites A|ATG at the 5′ end and T|TCG at the 3′ end. Consequently, the two vectors carrying GFP11 and GFP1-10 gene chimeras should be co-expressed in a single cell in order to determine protein targeting specificity to the organelle of interest.

### 3.4 Significance of high dual targeting frequency

The results obtained in this study strongly suggest that the number of dually targeted proteins is much higher than assumed so far. More than just increasing the list of such proteins, these findings also highlight the evolutionary conserved nature of organelle translocation machineries in plant cells. Even if some of these proteins are ‘mistargeted’ to the ‘wrong’ organelle, for example, due to protein overexpression, it shows the fundamental targeting properties of the respective transport signal. The reason of such widespread dual targeting is still open. On the one hand, it supports the hypothesis that dual targeting might be an evolutionary remnant (Staiger et al., 2009). On the other hand, the acquisition of dual targeting properties might still be ongoing process, which allows for the development of completely new biochemical pathways in an organelle (Martin 2010, Xu et al., 2013). It is, therefore, important to determine the targeting properties of protein transport signals in order to understand its physiological role in the plant cell. The *sa*split-GFP based approach and the PlaMiNGo toolkit developed here provide important tools to determine the targeting properties of a protein.

## 4. Materials and method

### 4.1 Molecular Cloning

#### 4.1.1 Generation of vectors for co-infiltration

The GFP1-10 and GFP11_x7_ fragments were amplified by PCR from plasmids pcDNA3.1-GFP (1-10) and pACUH-GFP11_x7_-mCherry-ß-tubulin (a gift from Bo Huang lab; Addgene#70218 and #70219) and cloned into pRT100mod-based vectors (Baudisch et al. 2014) either with digestion/ligation or with Restriction Free cloning (Bond and Naus, 2012). Primers for gene amplification are summarized in Suppl. Table 1. The gene sequence coding for the N-terminal 91 amino acids of TyrRS was amplified from a vector provided by E. Glaser and cloned accordingly (Baudisch et al. 2014). The above constructions, comprising promoter and terminator regions (*CaMV35S::Gene of interest:GFP1-10/GFP11_x7_::t35S*), were later sub-cloned into a *Golden Gate* compatible pLSU4GG binary vector with *Bsa*I restriction/ligation reaction (Erickson et al., 2017). *Golden Gate* cloning was performed in 15 µl reaction with following conditions: 2.5 units of T4 DNA ligase (Thermo Scientific), 5 units of *Bsa*I (Thermo Scientific), 1X BSA, 20 cycles of incubation at 37°C for 2 min and 16°C for 5 min, final deactivation and denaturation at 50°C for 10 min and 80°C for 10 min respectively.

#### 4.1.2 Construction of the PlaMiNGo toolkit

The modular cloning principle and DNA fragments of the Plant Parts I and II toolkits were used for vector construction (Weber et al., 2011; Engler et al., 2014; Gantner et al., 2018). The modules utilized for cloning of *Golden Gate*-based vectors are illustrated in Supplementary Figure 3. *Golden Gate* reactions were performed with 20 fmol of each DNA module with the following conditions: 2.5 units of T4 DNA ligase (Thermo Scientific), 5 units of *Bsa*I or *Bpi*I (New England Biolabs), 30 cycles of incubation at 37°C for 2 min and 16°C for 5 min, final denaturation at 80°C for 10 min. When required, the restriction/ligation reactions were subsequently supplemented with fresh ligation buffer and ligase for terminal ligation, and incubated for ≥ 3h at 16°C. Ligation mixtures were transformed into Dh10b or *ccd*B survival II cells (Thermo Scientific) and grown on plates with appropriate selective medium. PCR primers for Level 0 modules and gene sequences are summarized in Suppl. Table 2 and 3.

#### 4.1.3 Cloning of candidate proteins into PlaMiNGo vectors

The candidate targeting signals (TyrRS_1-91_, FNR_1-55_, mtRi_1-100_, ATPS_1-100_, GCS_1-100_, GAPDH_1-100_, Gtred_1-100_, RbcS_1-79_ and CoxIV_1-29_) were amplified from the corresponding cDNA templates (Nelson et al., 2007; Berglund et al., 2009; Baudisch et al., 2014) and cloned via standard *Golden Gate* reaction (see above) into PLaMiNGo vectors in exchange for a ccdB negative selection cassette. Overhangs of the fragment to be cloned were A|ATG at the 5′ end and T|TCG at the 3′ end. Two additional nucleotides were inserted in some of the fusions to maintain the reading frame resulting in an additional codon for an alanine residue.

#### 4.1.4 Cloning for complementation of the *immutans* phenotype

The gene sequence carrying mature PTOX_57-295_ was amplified from a cDNA clone provided by Steven Rodermel (Iowa State University, USA) and sub-cloned via restriction free cloning into pRT100mod-based vectors downstream of the gene sequences coding for the transport signals of either GCS_1-100_ or ATPS_1-100_. Additionally, the PTOX full-length gene and mature PTOX (lacking the transit peptide) were cloned into empty pRT100mod vectors under control of CaMV35 promoter and terminator. The candidate gene constructions with promoter and terminator (*CaMV35S::Candidate:PTOXmat::t35S*) were then cloned into a *Golden Gate* compatible pLSU4GG binary vector with *Bsa*I cut/ligation reaction.

### 4.2 Agrobacterium infiltration

The constructs were transformed into electro-competent cells of *Agrobacterium tumefaciens* GV3101 (pMP90) (Koncz and Schell, 1986). *Agrobacterium* infiltrations of 6 - 8 weeks old fully expanded *Nicotiana benthamiana* leaves were performed as described by Sharma et al. (2018a). For co-infiltration, each bacterial strain was adjusted to OD_600_ = 0.8 and mixed in a 1:1 ratio prior to infiltration.

### 4.3 Microscopy and imaging

Confocal laser scanning microscopy was carried out as described by Sharma et al. (2018a). For GFP/YFP dual channel imaging, the 493-518 (GFP) and 519-620 (YFP) filter ranges were used. When required, brightness and contrast of the images were equally adjusted for each image to avoid any discrepency in visualization of signal intensities.

### 4.4 Signal Quantification

For quantification of signal intensities, infiltration of all relevant constructs was carried out on different spots of the same leaf. At least three images from each infiltration spot were used for quantification. Quantification of the signals was performed with raw images using the Fiji program (Schindelin et al., 2012). For the purpose of quantification, image acquisition was done with the 20x objective in 7 to 8 Z-stacks covering the epidermal cell layer and later stacked to project the maximum intensities. The mean gray values of stacked images were calculated using the ‘Measure’ option of Fiji and further utilized for comparision of the fluorescence signal strenght in arbitrary units (A.U.).

### 4.5 Growth and transformation of immutans plants

The *Arabidopsis thaliana immutans* seeds (provided by Steven Rodermel, ISU) were germinated at 5 μmol m^−2^ s^−1^ lights and 8/16 h light-dark cycles. After three weeks of germination, the seedlings were transferred to 50 μmol m^−2^ s^−1^ lights. For induction of flowering, the plantlets were transferred to >150 μmol m^−2^ s^−1^ light and incubated at a 16/8 h light/dark cycle. Floral-dip transformation was performed as described (Davis et al. 2009). At least three independent T2 transgenic lines were selected in each case for the analysis of phenotype. After germination at >150 μmol m^−2^ s^−1^ light in a 8/16 h light/dark cycle, these plants were transferred to 16/8 h light/dark cycle. Pictures presented here are from 5 week old plants grown in >150-μmol m^−2^ s^−1^ light.

## Acknowledgement

We would like to thank Theresa Ilse for assistance in cloning, Elzbieta Glaser (Stockholm University, Sweden), Martin Schattat (Martin Luther University Halle-Wittenberg, Germany) and Bo Huang (University of California, San Francisco, USA; via Addgene) for kindly providing constructs and Steven Rodermel (Iowa State University, USA) for providing *immutans* mutants and the PTOX clone. MS was supported by a fellowship from the BRAVE project funded by the ERASMUS MUNDUS Action 2 program of the European Union.

## Supplementary File 1

Supplementary Figure 1

Confocal laser scanning microscopy of the lower epidermis of *Nicotiana benthamiana* infiltrated with *Agrobacterium tumefaciens* strain GV3101 carrying constructs encoding (a) FNR_1-55_/GFP1-10 or (b) mtRi1-55/GFP1-10 expressed under control of CaMV35S promoter. In both instances, no fluorescence signals were obtained. For further details, see the legend of Figure 2.

Supplementary Figure 2

Quantification of fluorescence signal intensity. Comparative analysis of signal intensities obtained in **(a)** plastids and **(b)** mitochondria with the use of co-infiltration and PlaMiNGo constructs. Image acquisition was done with 20-x objective and 2% of full laser power in several equally separated Z-stacks. Each bar represents the average pixel brightness from nine different digital images obtained from 2-3 independent experiments. Measure feature of Fiji was utilized for quantification of signals. (A.U.-Arbitrary units)

Supplementary Figure 3

Assembly of *Golden Gate*-based vectors and construction of the PlaMiNGo toolkit. The Modular Cloning system (Weber et al., 2011) and DNA modules from the Plant Parts I and II toolkits (Engler et al., 2014; Gantner et al., 2018) were used for vector assembly. All generated modules and vectors are summarized here. Level 0 modules were generated using gene specific primers (summarized in supplementary table 3). Oligonucleotides for GFP11_x1_ fragment were synthesized and hybridized before cloning. Five independent vector constructions (pTEI126, pTEI127, pTEI161, pTEI128, pTEI129) were generated in level I cloning steps. Two constructions namely pTEI176 and pTEI177 were generated additionally to extend the toolkit for the analysis of protein targeting to different sub-cellular compartments. The level I modules were then used to generate final vectors for analysis of protein targeting specificity to plastids (pLaNGo11-1, pLaNGo11-3, pLaNGo11-7) or mitochondria (pMiNGo11-1, pMiNGo11-3, pMiNGo11-7). These vectors carry a *ccd*B negative selection cassette flanked with *Bsa*I restriction site, which is replaced by cloning of the candidate transport signal.

Supplementary Figure 4

Subcellular localization of the candidate proteins **(a)** ATPS_1-100_/eYFP and **(b)** GCS_1-100_/eYFP, as determined by confocal laser scanning microscopy in leaf tissue of transgenic *Arabidopsis thaliana* lines. All images are maximum intensity projections of several single images representing the complete cell in z-axis. Overlay pictures of both the eYFP channel (displayed in yellow) and the chlorophyll channel (displayed in red) are shown. Separate images of the two channels are displayed at higher magnification in the middle panel (chlorophyll) and left panel (eYFP). A few chloroplasts are encircled with a while line for better display of signals. The scale bars correspond to 10 µm.

Supplementary Table 1-Primer Sequences (for cloning in pRT100 mod vector)

Supplementary Table 2-DNA sequence of (GFP1-10, YFP1-10, GFP11 tags and organelle transit peptides)

Supplementary Table 3-PCR Primers for cloning of Level 0 modules

