## Supplementary File 1 for "Determining targeting specificity of nuclear-encoded organelle proteins with the self-assembling split-fluorescent protein toolkit"

### Determining targeting specificity of nuclear-encoded organelle proteins with the self-assembling split fluorescent protein toolkit

Mayank Sharma, Carola Kretschmer, Christina Lampe, Johannes Stuttmann and Ralf Bernd Klösgen

**Supplementary Figure 1-**
Confocal laser scanning microscopy of the lower epidermis of *Nicotiana benthamiana* infiltrated with *Agrobacterium tumefaciens* strain GV3101 carrying constructs encoding (a) FNR1-55/GFP1-10 or (b) mtRi$_{1-55}$/GFP1-10 expressed under control of CaMV35S promoter. In both instances, no fluorescence signals were obtained. For further details, see the legend of Figure 2.

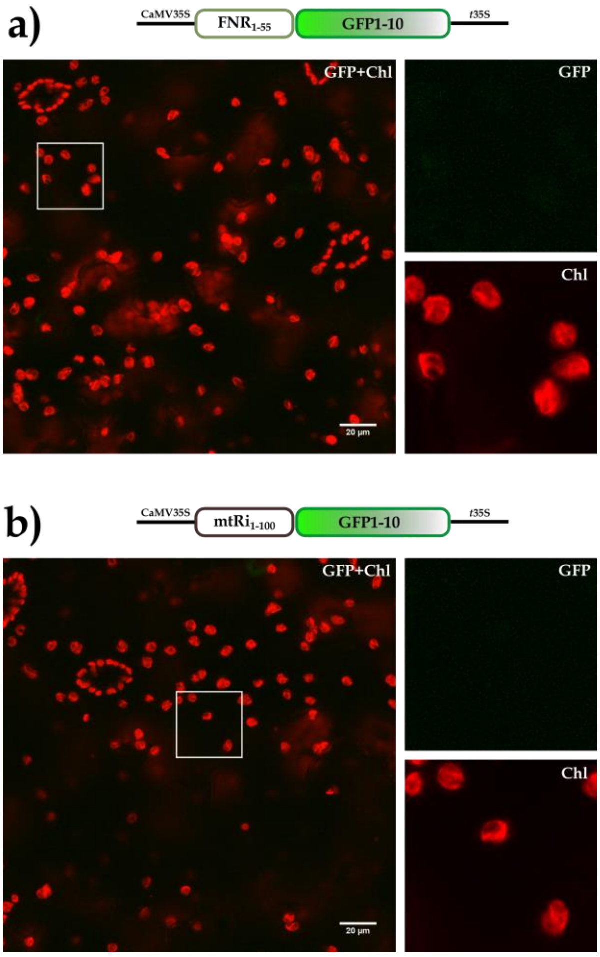

**Supplementary Figure 2-**

Quantification of fluorescence signal intensity. Comparative analysis of signal intensities obtained in **(a)** plastids and **(b)** mitochondria with the use of co-infiltration and PlaMiNGo constructs. Image acquisition was done with 20-x objective and 2% of full laser power in several equally separated Z-stacks. Each bar represents the average pixel brightness from nine different digital images obtained from 2-3 independent experiments. Measure feature of Fiji was utilized for quantification of signals. (A.U. - Arbitrary units)

**a)**

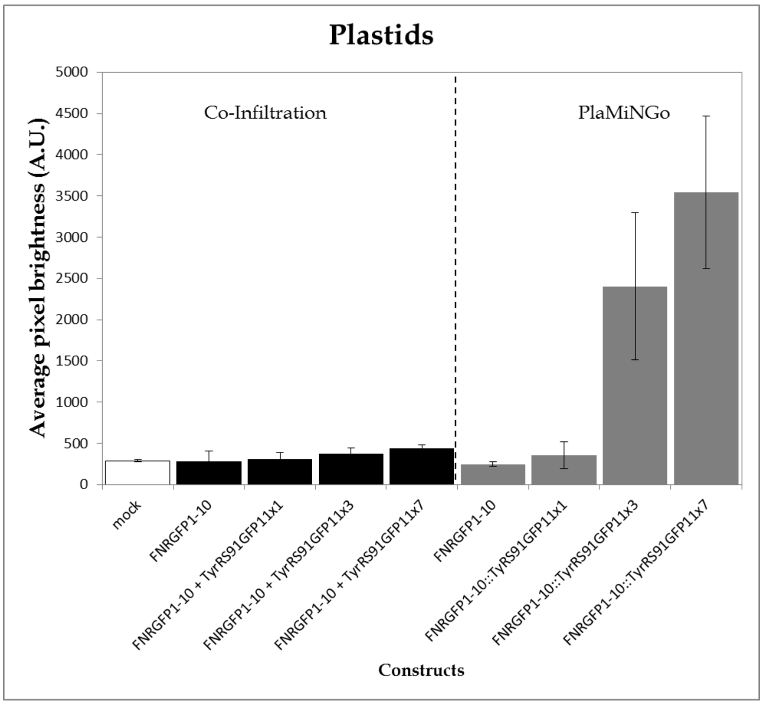


**b)**

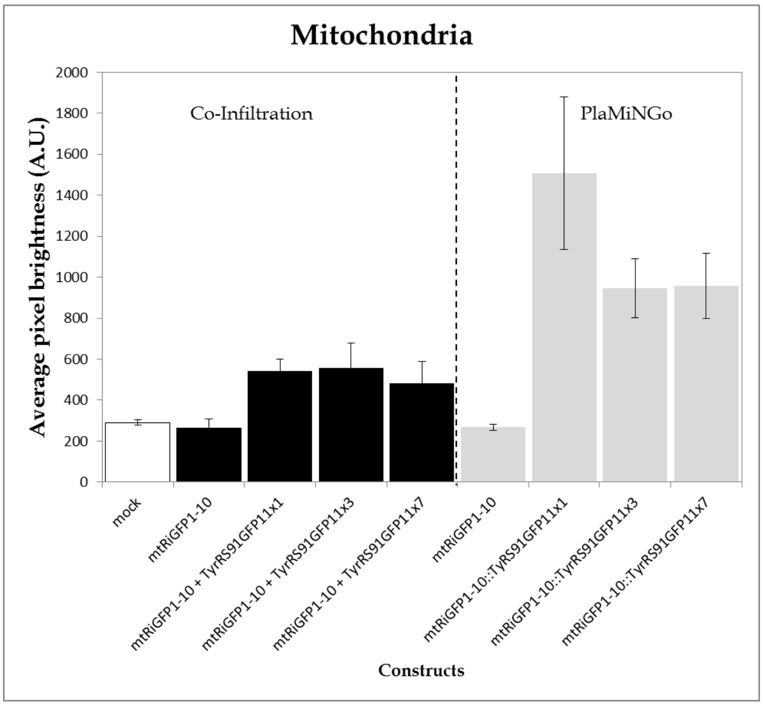

**Supplementary Figure 3-**

Assembly of Golden Gate-based vectors and construction of the PlaMiNGo toolkit. The Modular Cloning system (Weber et al., 2011) and DNA modules from the Plant Parts I and II toolkits (Engler et al., 2014; Gantner et al., 2018) were used for vector assembly. All generated modules and vectors are summarized here. Level 0 modules were generated using gene specific primers (summarized in supplementary table 3). Oligonucleotides for GFP11$_{x1}$ fragment were synthesized and hybridized before cloning. Five independent vector constructions (pTEI126, pTEI127, pTEI161, pTEI128, pTEI129) were generated in level I cloning steps. Two constructions namely pTEI176 and pTEI177 were generated additionally to extend the toolkit for the analysis of protein targeting to different sub-cellular compartments. The level I modules were then used to generate final vectors for analysis of protein targeting specificity to plastids (pLaNGo11-1, pLaNGo11-3, pLaNGo11-7) or mitochondria (pMiNGo11-1, pMiNGo11-3, pMiNGo11-7). These vectors carry a *ccd*B negative selection cassette flanked with *Bsa*I restriction site, which is replaced by cloning of the candidate transport signal.

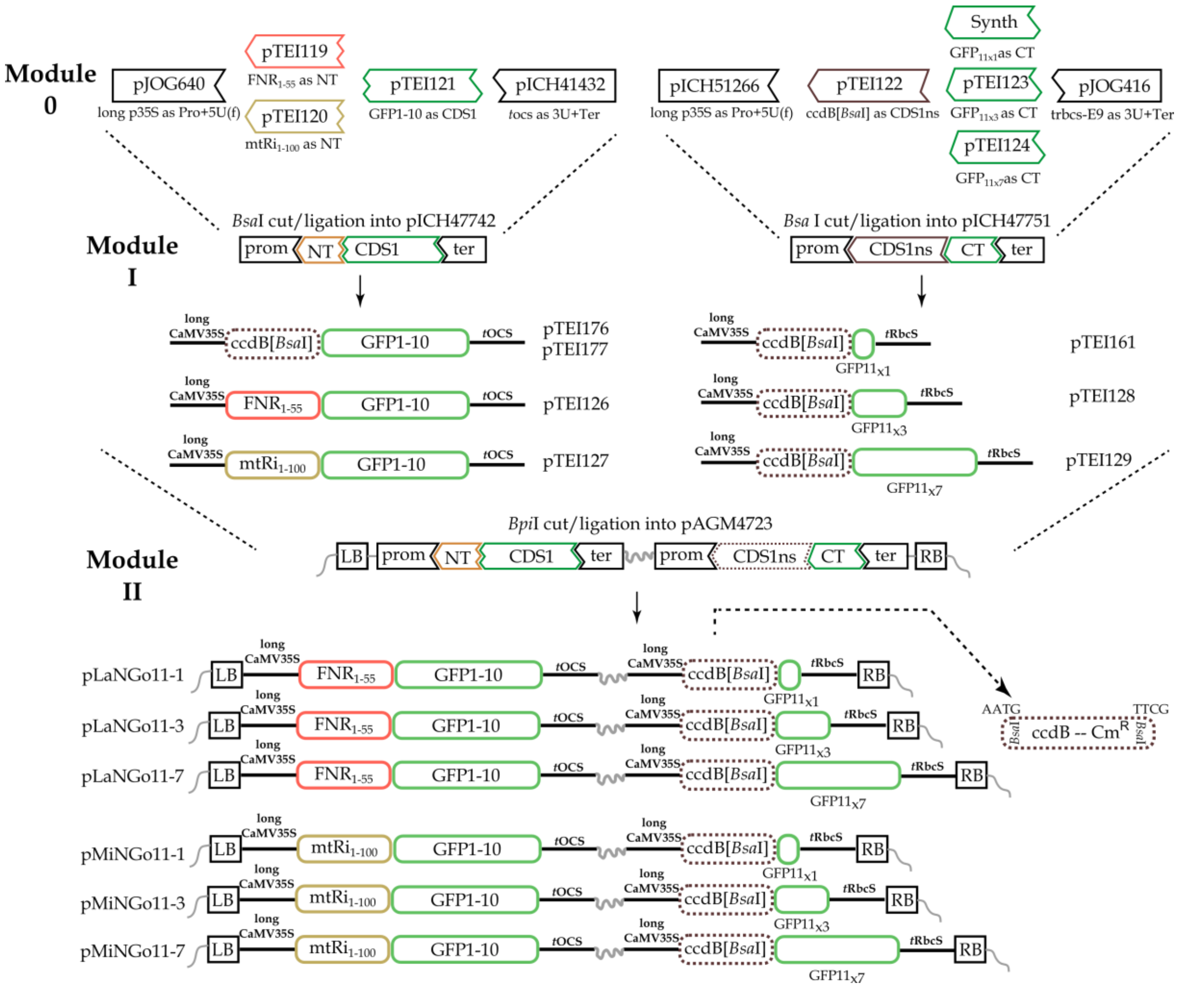

**Supplementary Figure 4-**

Subcellular localization of the candidate proteins **(a)** ATPS$_{1-100}$/eYFP and **(b)** GCS$_{1-100}$/eYFP, as determined by confocal laser scanning microscopy in leaf tissue of transgenic *Arabidopsis thaliana* lines. All images are maximum intensity projections of several single images representing the complete cell in z-axis. Overlay pictures of both the eYFP channel (displayed in yellow) and the chlorophyll channel (displayed in red) are shown. Separate images of the two channels are displayed at higher magnification in the middle panel (chlorophyll) and left panel (eYFP). A few chloroplasts are encircled with a while line for better display of signals. The scale bars correspond to 10 µm.

(a)

(b)

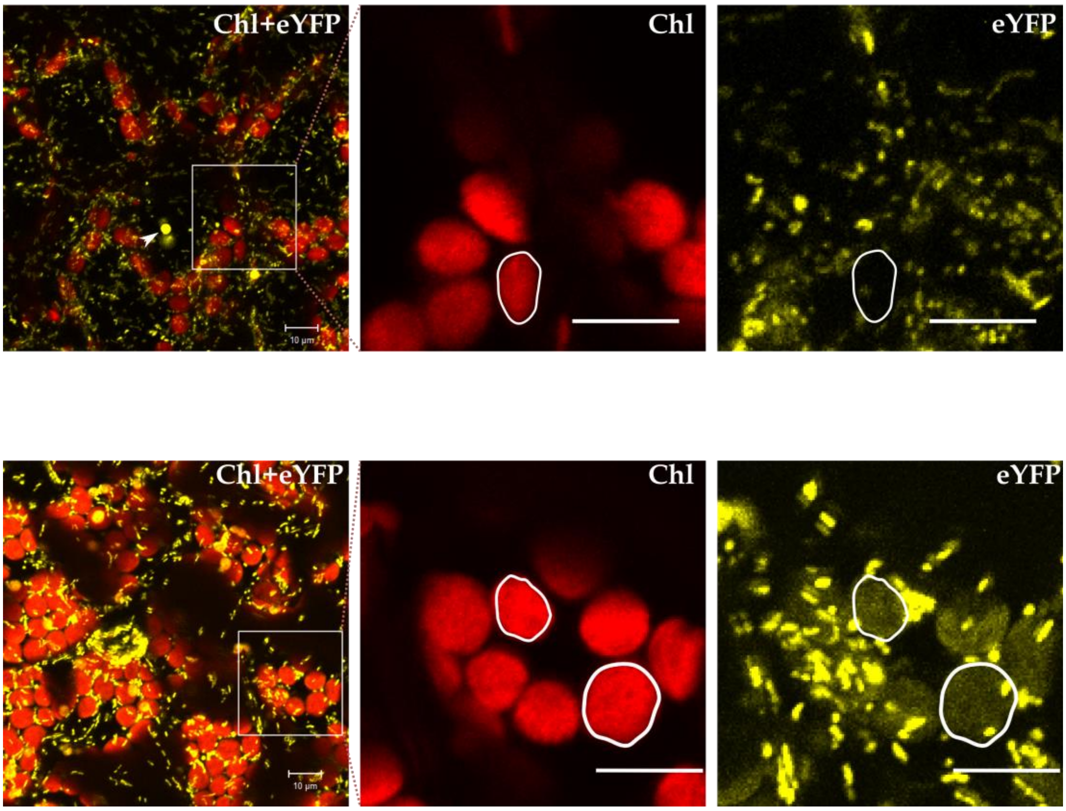

**Supplementary Table 1- Primer Sequences (for cloning in pRT100 mod vector)**

| Name | Oligonucleotide Sequence |
|---|---|
| GFP1-10 Forward | 5′ TATGGTCTCTCATGTCCAAAGGAGAAGAACTGTTTA 3′ |
| GFP1-10 Reverse | 5′ CCTCTAGACTAACTTCCGCCGCCACCT 3′ |
| GFP11 Forward | 5′ ATCCATGGGCGGCAAATTCATGCGTGACCACATGGT 3′ |
| GFP11x7 Reverse | 5′ ATTCTAGATTAGGTGATACCGGCAGCATTGAC 3′ |
| GFP11X3 Reverse | 5′ ATCTAGATTATCCGGTTATTCCGGCTGCATT 3′ |
| GFP11X1 Reverse | 5′ ATCTAGATTATGTAATCCCAGCAGCATTTAC 3′ |

Restriction site
Stop codon
Linker

**Supplementary Table 2- DNA sequence of (GFP1-10, YFP1-10, GFP11 tags and organelle transit peptides)**

| Name | Sequence (5' – 3') |
|---|---|
| GFP1-10 (Kamiyama et al., 2015) | ATGTCCAAAGGAGAAGAACTGTTTACCGGTGTTGTGCCAATTTTGGTTGAACTCGATGGT<br>GATGTCAACGGACATAAGTTCTCAGTGAGAGGCGAAGGAGAAGGTGACGCCACCATTGGA<br>AAATTGACTCTTAAATTCATCTGTACTACTGGTAAACTTCCTGTACCATGGCCGACTCTC<br>GTAACAACGCTTACGTACGGAGTTCAGTGCTTTTCGAGATACCCAGACCATATGAAAAGA<br>CATGACTTTTTTAAGTCGGCTATGCCTGAAGGTTACGTGCAAGAAAGAACAATTTCGTTC<br>AAAGATGATGGAAAATATAAAACTAGAGCAGTTGTTAAATTTGAAGGAGATACTTTGGTT<br>AACCGCATTGAACTGAAAGGAACAGATTTTAAAGAAGATGGTAATATTCTTGGACACAAA<br>CTCGAATACAATTTTAATAGTCATAACGTATACATCACTGCTGATAAGCAAAAGAACGGA<br>ATTAAAGCGAATTTCACAGTACGCCATAATGTAGAAGATGGCAGTGTTCAACTTGCCGAC<br>CATTACCAACAAAACACCCCTATTGGAGACGGTCCGGTACTTCTTCCTGATAATCACTAC<br>CTCTCAACACAAACAGTCCTGAGCAAAGATCCAAATGAAAAAGGAACAGGTGGCGGCGGA<br>AGTTAG |
| YFP1-10 | ATGTCCAAAGGAGAAGAACTGTTTACCGGTGTTGTGCCAATTTTGGTTGAACTCGATGGT<br>GATGTCAACGGACATAAGTTCTCAGTGAGAGGCGAAGGAGAAGGTGACGCCACCATTGGA<br>AAATTGACTCTTAAATTCATCTGTACTACTGGTAAACTTCCTGTACCATGGCCGACTCTC<br>GTAACAACGCTTACGTACGGAGTTCAGTGCTTTTCGAGATACCCAGACCATATGAAAAGA<br>CATGACTTTTTTAAGTCGGCTATGCCTGAAGGTTACGTGCAAGAAAGAACAATTTCGTTC<br>AAAGATGATGGAAAATATAAAACTAGAGCAGTTGTTAAATTTGAAGGAGATACTTTGGTT<br>AACCGCATTGAACTGAAAGGAACAGATTTTAAAGAAGATGGTAATATTCTTGGACACAAA<br>CTCGAATACAATTTTAATAGTCATAACGTATACATCACTGCTGATAAGCAAAAGAACGGA<br>ATTAAAGCGAATTTCACAGTACGCCATAATGTAGAAGATGGCAGTGTTCAACTTGCCGAC<br>CATTACCAACAAAACACCCCTATTGGAGACGGTCCGGTACTTCTTCCTGATAATCACTAC<br>CTCTCATATCAAACAGTCCTGAGCAAAGATCCAAATGAAAAAGGAACAGGTGGCGGCGGA<br>AGTTAG |
| GFP11x7 (Kamiyama et al., 2015) | ATGGGCGGCAAATTCATGCGTGACCACATGGTCCTTCATGAGTATGTAAATGCTGCTGGG<br>ATTACAGGTGGCTCTGGAGGTAGAGATCATATGGTTCTCCACGAATACGTTAACGCCGCA<br>GGCATCACTGGCGGTAGTGGAGGACGCGACCATATGGTACTACATGAATATGTCAATGCA<br>GCCGGAATAACCGGAGGGTCCGGAGGCCGGGATCACATGGTGCTGCATGAGTATGTGAAC<br>GCGGCGGGTATAACTGGTGGGTCGGGCGGACGAGATCATATGGTGCTTCACGAATACGTA<br>AACGCAGCTGGCATTACTGGCGGATCAGGTGGCAGGGATCACATGGTACTCCATGAGTAC<br>GTGAACGCTGCTGGAATCACAGGCGGTAGCGGCGGTCGGGACCATATGGTCCTGCACGAA<br>TATGTCAATGCTGCCGGTATCACCTAA |
| GFP11X3 | ATGGGCGGCAAATTCATGCGTGACCACATGGTCCTTCATGAGTATGTAAATGCTGCTGGG<br>ATTACAGGTGGCTCTGGAGGTAGAGATCATATGGTTCTCCACGAATACGTTAACGCCGCA<br>GGCATCACTGGCGGTAGTGGAGGACGCGACCATATGGTACTACATGAATATGTCAATGCA<br>GCCGGAATAACCGGATAA |

| | |
|---|---|
| GFP11X1 | ATGGGCGGCAAATTCATGCGTGACCACATGGTCCTTCATGAGTATGTAAATGCTGCTGGGATTACATAA |
| Mitochondria Marker (mtRi100) | ATGCTTCGAGTAGCAGGTAGAAGGCTTTCTTCTTCAGCCGCTAGATCTTCATCTACCTTCTTTACAAGAAGCTCTTTCACCGTTACCGATGATTCGTCTCCGGCAAGATCTCCTTCTCCGTCACTCACCTCTTCGTTTCTCGATCAAATCAGAGGTTTCTCATCTAATTCGGTTTCTCCCGCACATCAGTTGGGTTTAGTCTCAGATCTTCCAGCCACAGTGGCTGCTATTAAGAATCCCAGTTCAAAAATTGTATATGATGACTCCAACCATGAGCGTTATCCACCTGGAGATCCTAGC |
| Plastid Marker (FNRtp) | ATGACCACCGCTGTCACCGCCGCTGTTTCTTTCCCCTCTACCAAAACCACCTCTCTCTCCGCCCGAAGCTCCTCCGTCATTTCCCCTGACAAAATCAGCTACAAAAAGGTTCCTTTGTACTACAGGAATGTATCTGCAACTGGGAAAATGGGACCCATCAGGGCC |
| Dually targeted TyrRS91 (Elzbieta Glaser) | ATGGCATATGCAACAGGAATAACGTTTGCCTCAAGGAGTATTTTGCCTATTTGTTCCAGAACCTTCTTATCTCCCTTGCGTGTCGCTTCTTTACTCGTCTTTCCTGAGAAATCATCTGCAACTTTCTTCAGAAGGGTCCAAGTTCCTCACCTTTTTTCCACTTCTACTACTACTCTGTTCTCTTCTGTTAAGTGTTCAATTCATTCTACTTCATCTCCTGAGACAGAGAATCAAGCTGTTTTTCGCCCTAATGTAGTCGATATACTTGAAGAA |

Stop codon
Linker
Mutation for YFP

**Supplementary Table 3- PCR Primers for cloning of Level 0 modules**

| Module name | Oligonucleotide sequences (5' – 3') | Cloning vector |
|---|---|---|
| pTEI119 | tttgaagacatccATGACCACCGCTGTCACC | pAGM1276 |
| | tttgaagactaCATtgcGGCCCTGATGGGTCCCATTTTC | |
| pTEI120 | tttgaagacatccatGCTTCGAGTAGCAGGTAG | pAGM1276 |
| | tttgaagacatCATtgcGCTAGGATCTCCAGGTGGAT | |
| pTEI121 | tttgaagacataATGTCCAAAGGAGAAGAAC | pICH41308 |
| | tttgaagactaaagCTAACTTCCGCCGCCACCTG | |
| pTEI122 | tttgaagacatAATGtgagaccGACTGGCTGTGTATAAGGG | pAGM1287 |
| | tttgaagactaCGAAtgagaccTTGATCGGCACGTAAGAGG | |
| pTEI123 | tttgaagacattTCGATGGGCGGCAAATTCATGCG | pAGM1301 |
| | tttgaagactaaagcTTATCCGGTTATTCCGGC | |
| pTEI124 | tttgaagacattTCGATGGGCGGCAAATTCATGC (F1) | pAGM1301 |
| | tttgaagactaGaTCTCGTCCGCCCGACCC (R1) | |
| | tttgaagacatGAtCATATGGTGCTTCACGAATAC (F2) | |
| | tttgaagactaaagcTTAGGTGATACCGGCAGC (R2) | |
| pTEI125 | tttgaagacataATGTCCAAAGGAGAAGAAC (F1) | pICH41308 |
| | tttgaagacatGataTGAGAGGTAGTGATTATCAG (R1) | |
| | tttgaagactaaagCTAACTTCCGCCGCCACCTG (F2) | |
| | tttgaagacattatCAAACAGTCCTGAGCAAAG (R2) | |
| pTEI161* | tTCGATGGGCGGCAAATTCATGCGTGACCACATGGTCCTTCATGAGTATGTAAATGCTGCTGGGATTACATAA (F) | pICH47751 |
| | aagcTTATGTAATCCCAGCAGCATTTACATACTCATGAAGGACCATGTGGTCACGCATGAATTTGCCGCCCAT (R) | |

* For pTEI161, oligonucleotides were hybridized and cloned directly (GFP11$_{x1}$)

**Supplementary Video 1-**

Formation of aggregations with the use of multiple GFP11 tag. The PlaMiNGo vector carrying mtRi$_{1-100}$/GFP1-10 and TyrRS$_{1-91}$/GFP11x3 was infiltrated in *Nicotiana benthamiana* epidermal cells and analyzed via confocal microcopy. The time series presented here resulted from 60 consecutive digital images recorded at 6.25 s frame interval. A white arrow indicates the aggresome. *Time stamp*- Seconds, *Scale Bar*- 20 µM
